# Identification of potential carboxylic acid-containing drug candidate to design novel competitive NDM inhibitors: An *in-silico* approach comprising combined virtual screening and molecular dynamics simulation

**DOI:** 10.1101/2021.07.05.451101

**Authors:** Karthick Vasudevan, Soumya Basu, Amala Arumugam, Aniket Naha, Sudha Ramaiah, Anand Anbarasu, Balaji Veeraraghavan

**Author notes:** **Corresponding Author details: 1. Dr Anand Anbarasu, Email:**, **2. Dr Balaji Veeraraghavan, Email:**. Karthick Vasudevan and Soumya Basu contributed equally to this manuscript.

## Abstract

Metallo-β-lactamases (MBLs) producing bacteria especially the ones with New Delhi metallo-beta-lactamase-1 (NDM-1) and its variants can potentially hydrolyse all the major β-lactam antibiotics, ultimately escalating anti-microbial resistance world-wide. There is a dearth of approved inhibitors to combat NDM and other MBLs producing bacteria. Hence we focussed to find novel inhibitor(s) *in-silico* which can potentially suppress the activity of NDM/ MBLs. 2400 compounds were virtually screened to identify a promising carboxylic acid-containing compound (CID-53986787) analogous to NDM antagonist Captopril. Our lead compound can bind adjacent to the active site zinc ions (Zn1 and Zn2) in all highly resistant NDM variants. CID-53986787 possesses ~5-8% higher binding affinity than Captopril, exhibiting molecular interactions with crucial residues that can destabilize the hydrolytic activity of NDM. CID-53986787 was virtually evaluated to ascertain its safe pharmacological/ toxicity profile. Molecular dynamics simulation studies elucidated its stable interaction with the target protein (NDM-1).

## INTRODUCTION

Antibiotic resistance is the growing concern and major threat to human health. β-lactamases are important class of enzymes present in bacteria that can efficiently hydrolyse the β-lactam ring thereby rendering the antibiotics ineffective. β-lactamases are classified into four major classes. Classes A, C, and D encompasses serine residue in its active site and promotes catalysis [1,2]; whereas, class B possess two zinc metal ions for hydrolysis of antibiotics which therefore referred to as metallo-β-lactamases (MBLs) [3]. His120, His122 and His189 are Zn1 co-ordinating residues, whereas Asp124, Cys208 and His250 are Zn2 co-ordinating amino acid residues [4]. The MBL encoding genes can spread between bacteria through horizontal transfer mechanism, thereby converting non-resistant bacteria to resistant bacteria. These enzymes can even potentially degrade carbapenems, which is considered as last line of defence for the treatment against pathogenic bacteria [5].

Carbapenem resistance (CR) in Gram-negative organisms is increasing over the years across the globe, reported with high morbidity and mortality rates [6]. Molecular mechanisms of CR are due to production of acquired carbapenemases that degrades carbapenem and/or chromosomal mediated loss of porins as well as over-expression of efflux pumps. Carbapenemase mediated resistance is a major concern, as they rapidly disseminate through mobile genetic elements. Very few agents like Colistin, Tigecycline and Fosfomycin are being used for CR infections. Due to the limited therapeutic options, inhibitor molecules are discovered to be partnered with a suitable beta lactam agent, in turn to act against several classes of beta lactamases including carbapenemases.

Newer Beta Lactam/Beta Lactamase Inhibitors (BL/BLIs) agents includes: Ceftazidime/ Avibactam, Ceftolozane/ Tazobactam, Aztreonam/ Avibactam, Imipenem/ Relebactam, Meropenem/ Vaborbactam, Cefepime/ Zidebactam and Meropenem/ Nacubactam [7]. These agents work against specific type of carbapenemases with majority being active against Class-A carbapenemases (KPC). However, NDM and Class-D oxacillinases (Oxa-48 like) are endemic in Indian settings [8] with limited utility of these newer agents. The *bla_NDM-1_* gene and its variants is located on plasmids harbouring multiple resistant determinants, thereby conferring extensive drug resistance, leaving only a few or no therapeutic options that require rapid development of inhibitors [9–11]. As NDM is highly endemic across Indian subcontinent [12], it is the need of the hour to have active MBL inhibitors for the management of pathogen infestations. Our research group has extensively worked in the field *in-silico* prediction of antimicrobial therapeutics and structural biology to understand the protein-ligand interaction against pathogenic microbes [13–15].

D-Captopril is a carboxylic acid based compound that found to be having antagonistic effect on the activity of NDM-1 [16]. Thus, the screening of D-Captopril analogues using virtual tools and trusted databases [17] could shed light on the inhibitor design and effective treatment of pathogenic bacteria carrying NDM-1 and its variants.

## MATERIALS AND METHODS

### Datasets

The three-dimensional structures of NDM-1 (5ZGE) and NDM-5 (4TZE) were retrieved from protein databank [18,19]. Meanwhile, 3D structure of NDM-7 is modelled using Swiss Model and the most appropriate model was selected on the basis of QMEAN value [20]. The lead compounds, ≥ 80% similar to D-captopril were taken from PubChem database, converted to PDB format and checked to avoid sequence breaks or other anomalies. The target proteins were energy minimized prior to docking analysis [21,22].

### Receptor based Virtual screening and Bioavailability analysis

Receptor based virtual screening [23] is computational alternative for traditional drug screening. Recently, this approach has been widely utilized for drug candidate identification in many pharmaceutical and research organisations. The idea of virtual screening is to reduce the duration of lead discovery, prioritize the compounds based on its target specific binding and rapid development of drug candidates with minimal cost [24]. In this study, we utilized PubChem database to screen carboxylic acid-containing D-captopril analogues with similarity index of 80% and prioritized the lead candidates based on binding affinity. Absorption, distribution, metabolism, elimination and toxicity (ADME/Tox) are the crucial factors that will determine the clinical approval of a drug candidate. Most of these compounds fail at pre-clinical/clinical phase due to its higher toxicity and reduced efficacy [25]. Lipinski Rule of Five is the most promising methodology used in characterizing the drug candidates based on its ADME properties [26]. In the present study, these ADME related properties of the screened compound were determined using MOLINSPIRATION program (2008).

### Molecular Docking analysis

We further evaluated the binding affinity of the NDM variants and lead compounds with molecular docking analysis using AutoDock Vina [27,28]. Docking was performed to illustrate the binding mechanism of the ligand to the target binding site and determine its crucial interactions. Repeated docking was employed to generate more accurate binding affinity and to eliminate false positive results. Grid box with appropriate dimensions was used to encompass the entire active site residues. Furthermore, the intermolecular interactions of the complexes were analysed to determine the crucial residues of the target that can contribute to the enzymatic activity.

### Prediction of toxicity risks

Toxicity and poor pharmacokinetics are the major hindrance in development of new drugs. Approval of drug requires drug candidates possessing good pharmacokinetic properties and should satisfy all drug-like properties and failure which should be eliminated in the early stages of drug development [25]. We utilized Osiris property explorer (https://www.organic-chemistry.org/prog/peo/) and Protox web server, to evaluate the toxicity and side effects of the screened lead compounds [29]. Further, the predicted compound stability was evaluated and compared with D-captopril using Molecular dynamics simulations.

### Molecular dynamics simulation

The Simulations were carried out using GROMACS package to determine the stability of the bound complexes throughout the simulation time. GROMOS43a1 force field is utilized for all the simulations and PRODRG server is used for generation of lead molecules topologies. The complexes were then solvated in a cubic box of 0.9nm with SPC water model and sodium counter ions were added to neutralize the entire system [21,22]. Steepest descent algorithm was utilized for energy minimization to bring the system into energetically favourable condition and to relax the bond lengths and angles. In addition, the entire system was equilibrated using position-restrained dynamics simulation in terms of pressure and temperature (NPT and NVT). Finally, 50ns molecular dynamics simulations ware carried out at optimal pressure and temperature. Particle-mesh Ewald algorithm were used for long-range electrostatic interactions [30,31]. Each snapshot were taken at the rate of 1ps and stored in the trajectory. The deviation and hydrogen bonding strength of the complexes were calculated using RMSD and H-bond analysis. The binding energy between lead candidate and target is analysed using MM-PBSA approach [32].

## RESULTS AND DISCUSSIONS

### Receptor based Virtual screening and Bioavailability analysis

Around 2400 compounds with ≥80% similar to D-captopril were screened from the PubChem database. Molecular docking was carried between each lead compound and NDM-1. To increase the accuracy and reliability of the result, each docking calculation was repeated and mean was calculated. Top ranked compounds possessing higher binding affinity were selected for further analysis.

For acceptable drug candidate, it should not have molecular weight of more than 500, the LogP and hydrogen bond donors is not more than 5 and the hydrogen bond acceptor is not more than 10. These molecular properties of the screened lead compounds were calculated using the MOLINSPIRATION according to Lipinski rule of five [26]. This analysis depicted that all the screened compounds were in acceptable criteria [**Table 1**].

**Table 1:**
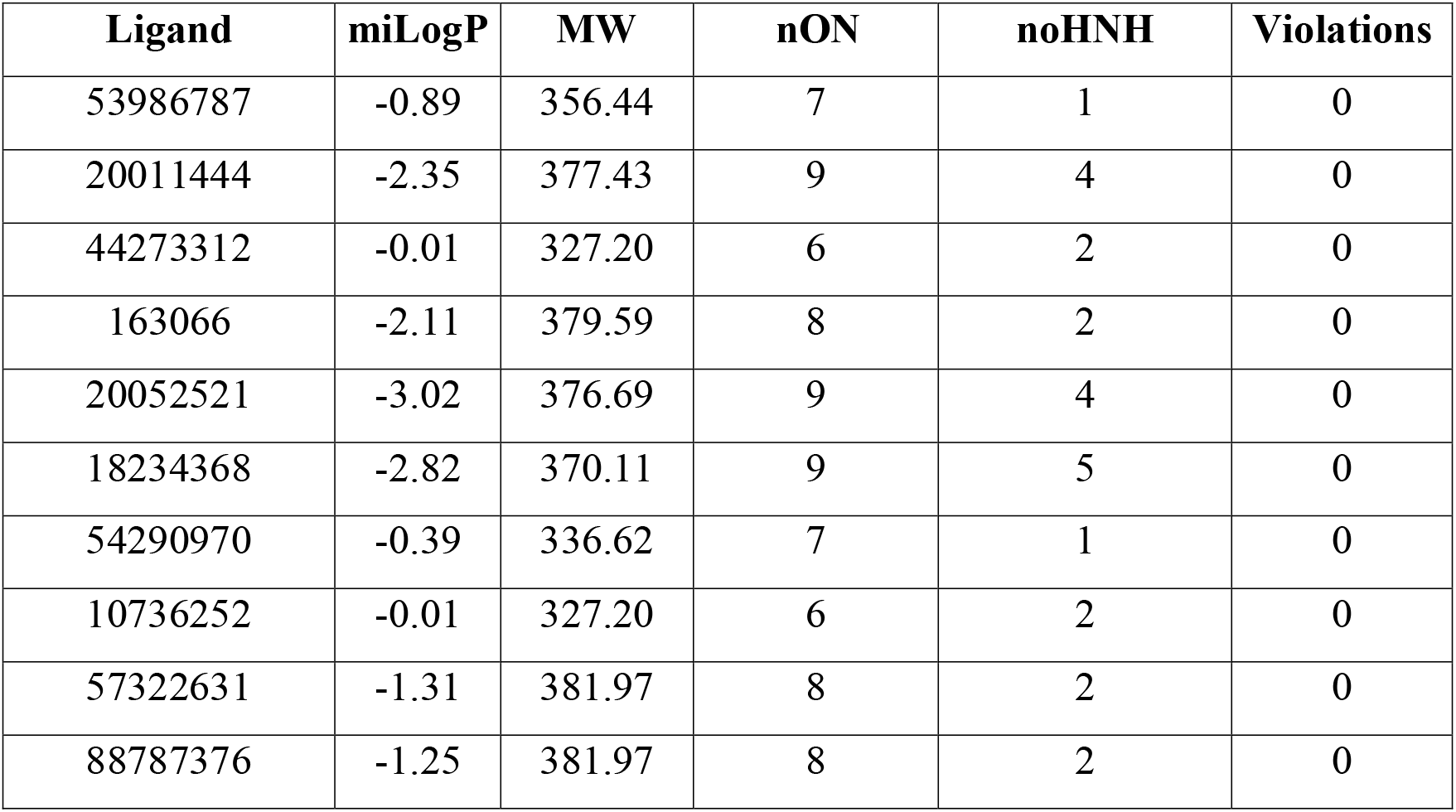
Calculations of molecular properties of lead compound using Molinspiration.

### Molecular Docking based virtual screening

We further docked the screened compounds with NDM-1 and other predominant variants that can lead to antibiotic resistance. The result showed that the 53986787 (compound 1) found to have higher binding affinity with NDM-1, NDM-5 and NDM-7 variants when compared to D-captopril and other lead compounds [**Table 2**]. In addition, Discovery studio and PyMol [33,34] used to visualize the intermolecular interactions present in the docked complex structures. The interacting pattern and its respective molecular function of the residues are shown in the **Figure 1** and **Figure 2**.

**Table 2:**
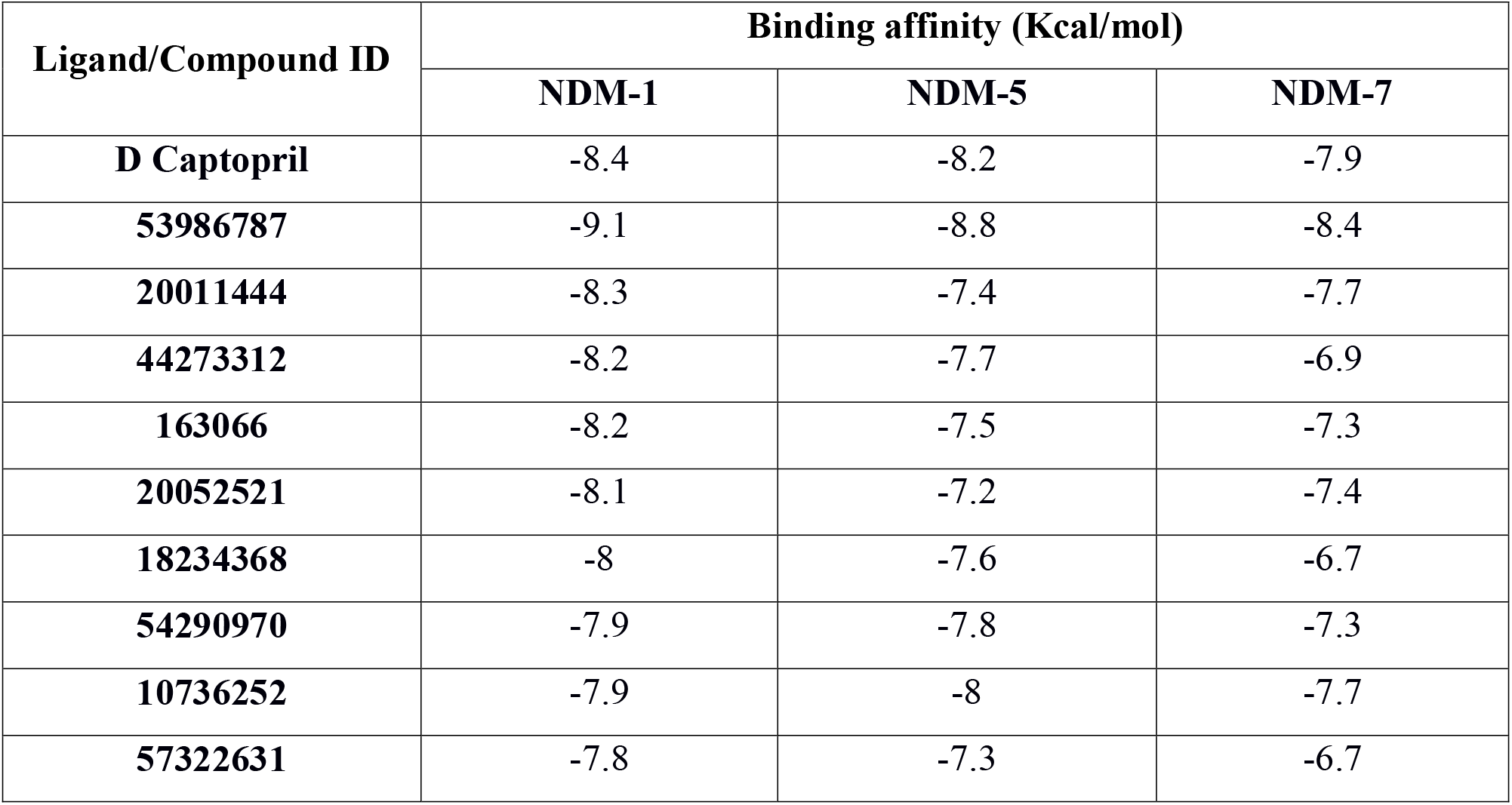
Molecular docking analysis between NDM-1 and other important variants with lead compounds

| Ligand/Compound ID | Binding affinity (Kcal/mol) |  |  |
| --- | --- | --- | --- |
|  | NDM-1 | NDM-5 | NDM-7 |
| D Captopril | -8.4 | -8.2 | -7.9 |
| 53986787 | -9.1 | -8.8 | -8.4 |
| 20011444 | -8.3 | -7.4 | -7.7 |
| 44273312 | -8.2 | -7.7 | -6.9 |
| 163066 | -8.2 | -7.5 | -7.3 |
| 20052521 | -8.1 | -7.2 | -7.4 |
| 18234368 | -8 | -7.6 | -6.7 |
| 54290970 | -7.9 | -7.8 | -7.3 |
| 10736252 | -7.9 | -8 | -7.7 |
| 57322631 | -7.8 | -7.3 | -6.7 |

**Figure 1:**
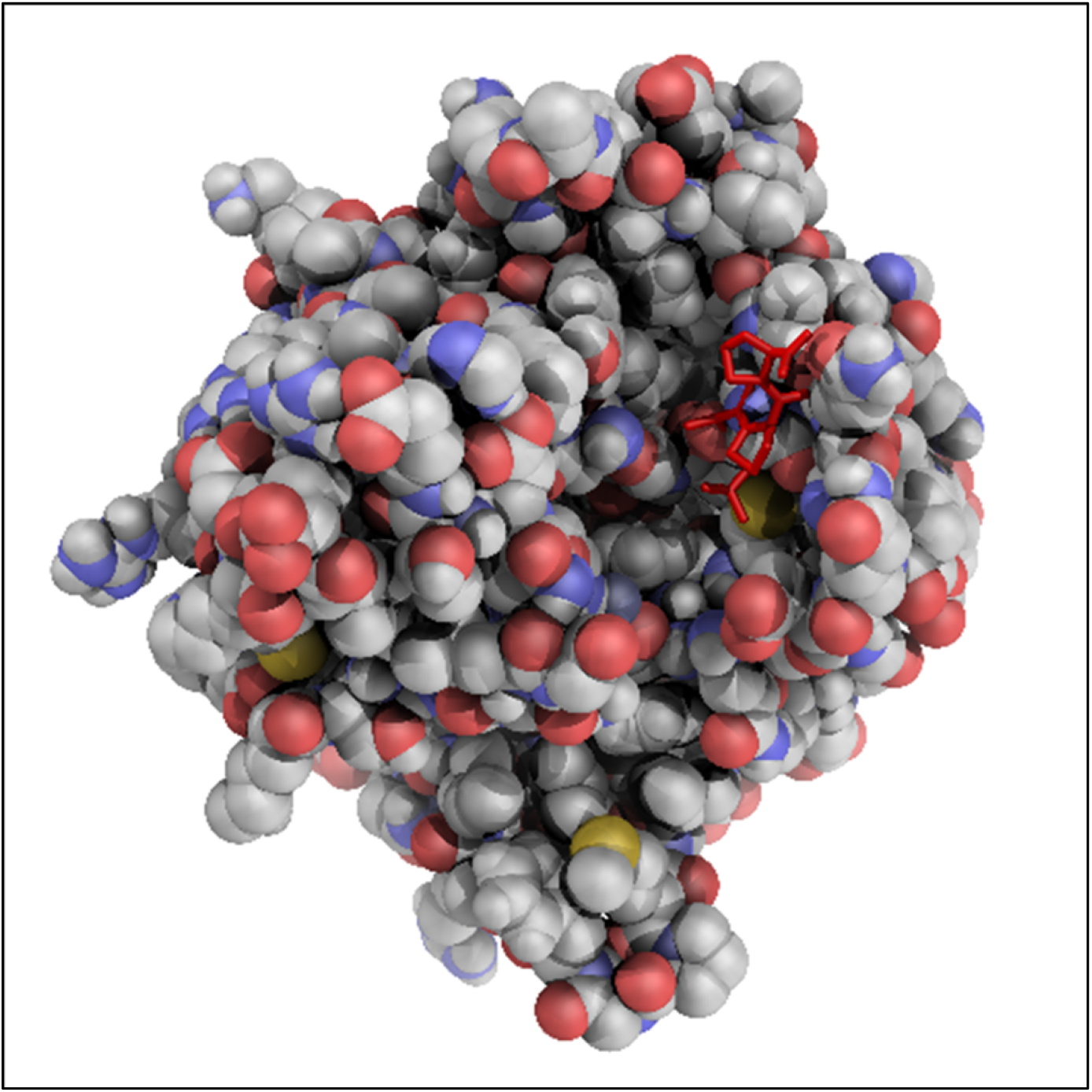
The docked complex of NDM-1 and Compound-1 depicting the binding of the compound in the active site of the target

**Figure 2:**
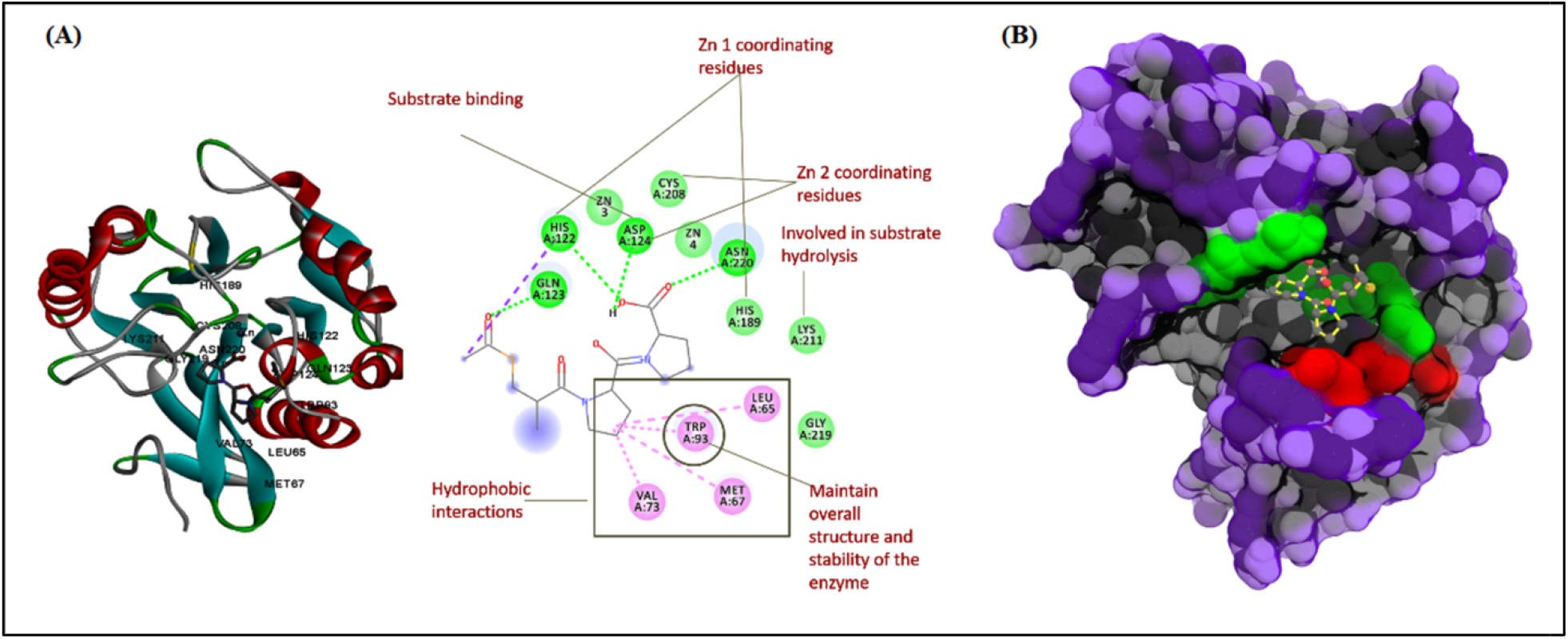
Crucial interacting residues between the compound 1 and NDM-1 and their respective molecular function in maintaining the stability of the enzyme (A) and surface view representing the binding mode of Compound-1 with both active site and non-active site residue

The complex exhibits crucial interaction with both Zn1 (His-122, His-189) and Zn2 co-ordinating residues (CYS-208, ASP-124). In specific, conventional hydrogen bond interaction was found with HIS-122, ASP-124, ASN-220 and GLN-123. In addition, van der Walls interaction was found with LYS-211, GLY-219, HIS-189 and CYS-208. Further, the compound 1 exhibited hydrophobic interactions with LEU-65, MET-67 and VAL-73 and also with non-active site residue TRP-93 which is related to maintenance of structure and stability of the enzyme [35] [**Figure 2**]. This analysis indicated that the compound 1 can efficiently bind to the active site of the NDM-1by competitive binding mechanism and prevent the hydrolytic activity of the enzyme.

### Prediction of toxicity risks

*In silico* prediction of LD50 can efficiently reduce the failure of drug in pre-clinical stage. It also serves in reducing the time and cost by ranking the compounds on the basis of safety profiles prior to animal experiments. The predicted LD50 value of the screened lead compound is shown to be 5000 mg/kg and also found to be non-mutagenic, non-tumorigenic and no reproductive or irritation side effects with drug-likeness and drug score of 3.26 and 0.896 respectively. The positive drug-likeness indicates the screened compound contains fragments present in approved commercial drugs. This analysis show that this predicted compound exhibits better safety profiles and less likely to cause any side effects.

### Molecular dynamics simulation

Although docking based virtual screening is very essential in prioritizing the lead compounds on the basis of binding affinity, it cannot reveal the protein dynamics after binding of the ligand. Molecular dynamics can efficiently decipher the binding dynamics and ligand-induced conformational changes. In the present study, molecular dynamic simulation was carried out by GROMACS package 5.1.4 adopting the GROMOS43a1 force field parameters which aimed to simulate the ligand induced potential conformational movements of the docked complexes.

### Distance calculation

Non-active site residue (TRP-93) plays an important role in maintaining the stability of NDM-1 [36]. The distance calculated between TRP-93 and lead compounds also revealed that Compound-1 exhibited closer interaction with D-captopril [**Figure 3(A)**]. From this analysis, it is evident that these crucial active and non-active site interactions will be helpful in stable binding of the Compound-1 to NDM-1. Similarly, LYS-211 is the important active site residue that helps in substrate recognition and hydrolysis [35]. The distance calculated between LYS-211 and lead compounds revealed that Compound-1 exhibits closer interaction with D-captopril [**Figure 3(B)**].

**Figure 3:**
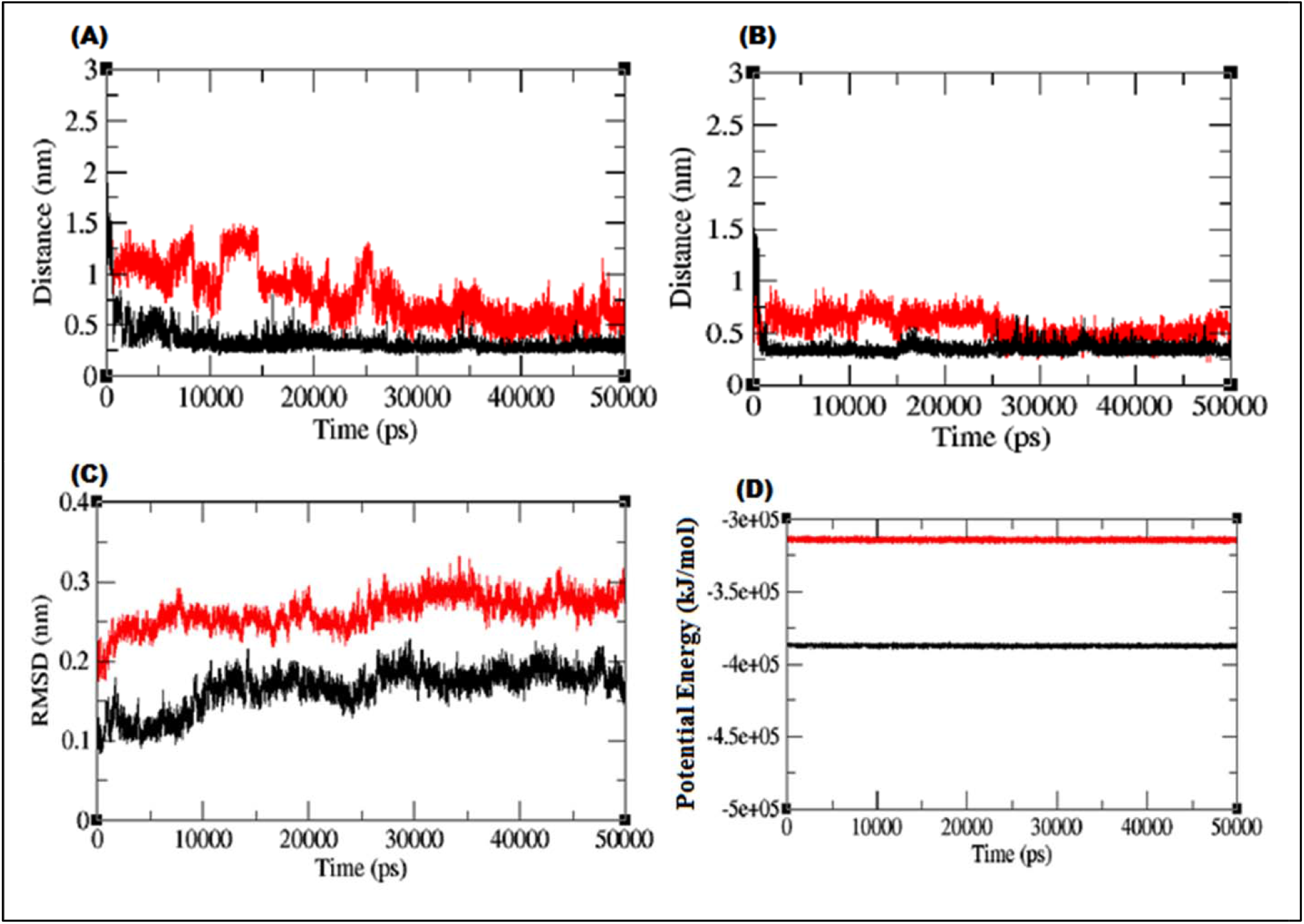
Molecular Dynamic Simulation: **(A)** The distance between Trp-93 and NDM-1-compound1 (black) and between Trp-93 and NDM-1-captopril (red) **(B)** The distance between LYS-211 and NDM-1-compound1 (black) and between Lys-211 and NDM-1-captopril (red) **(C)** RMSD of NDM1-compound1 (black) and NDM1-captopril (red) **(D)** Potential energy of NDM1-Compound-1 (black) and NDM1-Captopril (red)

### RMSD analysis and Potential energy

The root mean square deviation (RMSD) analysis showed that the NDM-1-compound displayed an RMSD of ~0.15 at 50 ns into the simulation and whereas the NDM-1-captopril interaction displayed an RMSD of ~0.3 at the end of the simulation suggesting that screened compound show more stability than D-captopril [**Figure 3(C)**]. In addition, potential energy analysis revealed that the both complexes were energetically stabilized throughout the simulation process [**Figure 3(D)**].

### Hydrogen bond analysis

The number of hydrogen bond directly correlates with binding strength of the ligand to the protein. From NHBOND analysis, the Compound-1 possesses 4-5 hydrogen bonds, whereas D-captopril exhibit only 3 hydrogen bonds throughout the simulation time. This binding strength is further evaluated with MM-PBSA and molecular dynamics based docking analysis.

### Molecular dynamics-based docking analysis

The trajectory was extracted at different conformations at the interval of 5ns. The docking was performed at each of the conformation with both the compounds. The result indicated the Compound-1 possessed significantly higher binding affinity at each of the conformation [**Figure 4**].

**Figure 4:**
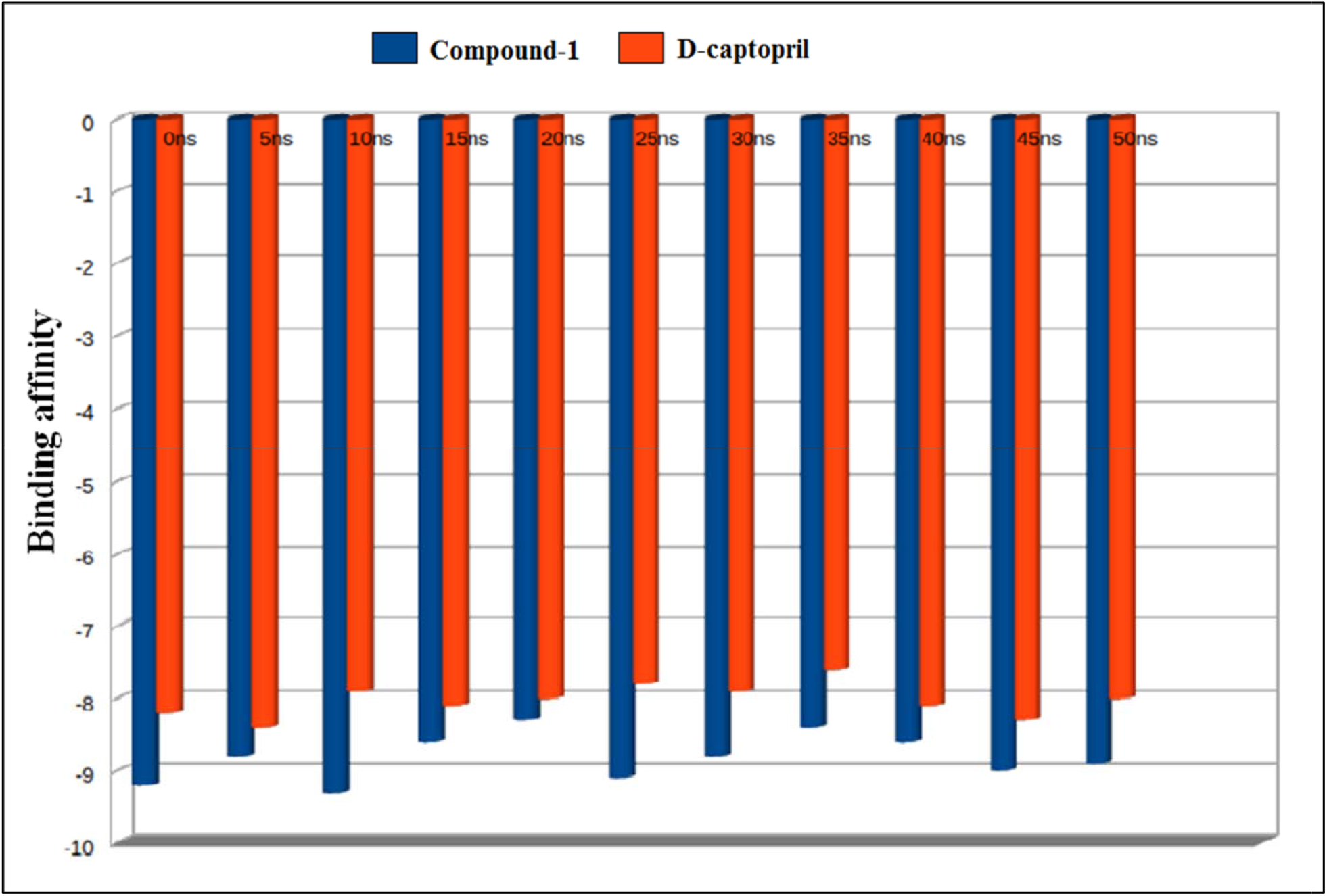
Molecular dynamics based docking analysis between NDM-1-compound1 and NDM-1-captopril (orange) complexes

### MM-PBSA analysis

To further evaluate the binding affinity of lead compounds, MM-PBSA analysis was used to calculate the binding affinity between protein and ligand from each of the conformation of protein during molecular dynamics simulation. It is clear from the analysis that the lead compound exhibited higher binding affinity throughout the simulation time [**Figure 5**], which illustrates the competitive and efficient binding of the Compound-1 to the target.

**Figure 5:**
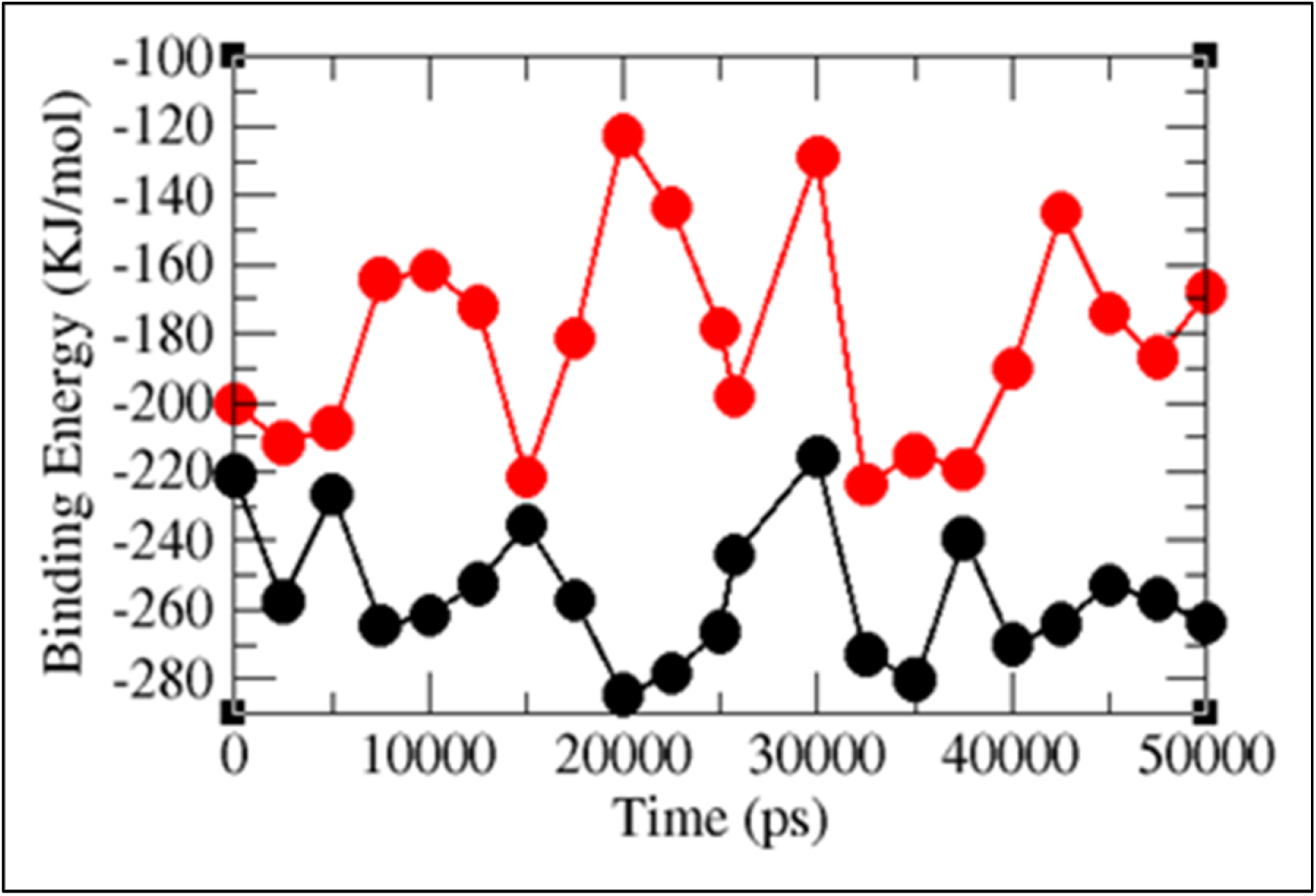
MMPBSA based binding energy calculation between NDM-1-compound1 (black) and NDM-1-captopril (red) complexes throughout the simulation time

## CONCLUSIONS

Resistance to antibiotics is the major threat to human health that causes failure of therapeutics. NDM producing MBLs are highly resistant to major class of antibiotics which necessitates the rapid development of new inhibitors that can competitively bind to the target thereby promoting the efficacy of antibiotic therapy. The docking analysis depicted that Compound-1 have crucial interactions with most of the key residues which is required for NDM hydrolytic activity. Finally, molecular dynamics simulation results also validated the effective binding of Compound-1 with the target. In addition, predicted oral toxicity and other physicochemical properties showed that CID 53986787 (Compound-1) can be safer and potential drug candidate for the future development of NDM inhibitors.

## CONFLICTS OF INTEREST

The authors declare that they have no conflicts of interest.

## FUNDING

The authors gratefully acknowledge Indian Council of Medical Research (ICMR), Govt. of India for the research grant **IRIS ID: 2019-0810** to carry out the present research.

## ACKNOWLEDGMENTS

The authors would like to thank the management of the Department of Clinical Microbiology, CMC-Vellore and School of Bio-Sciences and Technology, VIT-Vellore, for providing the necessary facilities to carry out this research work. Soumya Basu and Aniket Naha sincerely thank ICMR for his research fellowship.

## AUTHOR CONTRIBUTIONS

**Karthick Vasudevan** and **Soumya Basu:** Data Curation, *In-silico* experimentation and analysis, Writing-original draft; **Amala Arumugam:** *In-silico* experimentation and analysis; **Aniket Naha:** Literature Survey, Data analysis; **Sudha Ramaiah:** Validation of methodology and manuscript review; **Anand Anbarasu** and **Balaji Veeraraghavan:** Conceptualisation, Project Administration, Funding Acquisition, Manuscript review

## Notes

### Competing Interest Statement

The authors have declared no competing interest.

